# *StPOPA*, encoding an anionic peroxidase in *Solanum tuberosum*, enhances resistance against *Phytophthora infestans*

**DOI:** 10.1101/431346

**Authors:** Yu Yang, Rui Jiang, Hongyang Wang, Zhendong Tian, Conghua Xie

**Author notes:** Corresponding author (C. Xie).

## Abstract

Potato late blight, caused by *Phytophthora infestans*, is one of the major threats affecting the quality and output of potato all over the world. Reactive oxygen species (ROS) acted as a signal molecule to transmit signals in plants at the early stage of disease infection, and it could induce disease resistance of the plant, including potato late blight. Anionic peroxidases in many plants were reported to be involved in defense to disease. However, limited information about anionic peroxidase genes is available for the potato. Here, we reported that the expression of the *StPOPA*, a gene encoding a suberization-associated anionic peroxidase, was associated with resistance in potato against *P. infestans.* The *StPOPA* gene was induced by *P. infestans* infection, mechanical damage, jasmonic acid and ethylene treatment. Overexpression of the *StPOPA* gene in potato enhanced the resistance against *P. infestans* via promoting the accumulation of callose in the cell wall and ROS in the cytoplasm, which restricted the infection and spreading of the disease possibly by purposeful programmed cell death. Taken together, our results suggested that the *StPOPA* gene contributed to potato immunity against *P. infestans* and this gene could be used for the genetic improvement of resistance against potato late blight.

## Introduction

Potato (*Solanum tuberosum*) is the fourth largest food crop in the world. However, late blight, caused by the oomycete pathogen *Phytophthora infestans*, is seriously affecting potato production all over the world (Jiang et al., 2018). Unfortunately, no effective ways to control late blight have yet been found except for frequent use of chemicals. The most efficient strategy to prevent the disease is to develop crop varieties with durable and broad-spectrum resistance (Dangl et al., 2013; Fukuoka et al., 2009).

Plants are attacked by bacteria, viruses, fungi, oomycetes and other pathogenic microorganisms in their lives and could defend themselves via multiple resistance mechanisms (Giraldo and Valent, 2013). Despite the lack of a cellular immune system, plants share with animals an innate immune system (Li et al., 2017). There are two major innate immune responses in plants: the pathogen-associated molecular pattern (PAMP)-triggered immunity (PTI) (Boller and He, 2009) and the effector-triggered immunity (ETI) (Jones and Dangl, 2006). The PTI and ETI phases of plant immunity may be spatiotemporally distinct, but they are both intimately related to the reactive oxygen species (ROS) burst (Grant and Loake, 2000; Kadota et al., 2014; Nürnberger et al., 2004; Torres et al., 2006).

As one part of the plant defense response mechanism, the outbreak of reactive oxygen species is closely related to the occurrence of the anaphylactic reaction and programmed cell death (Lamb and Dixon, 1997) and production of ROS in plant cells marks the successful recognition of plant pathogens and activation of plant defenses (Torres, 2010). Doke first demonstrated pathogen-induced apoplastic ROS production in potato tuber tissues (Doke, 1983) and indeed, ROS was reported to act as signal transduction molecules in plant defense response (Frederickson Matika and Loake, 2013; Lehmann et al., 2015; Mittler et al., 2011; Torres, 2010).

Peroxidases (EC1.11.1.7) can catalyze the redox reaction of hydrogen peroxide with various inorganic hydrogen or organic hydrogen donors (Jouili et al., 2011). Welinder divided peroxidases superfamily into three categories: Class I, Class II and Class III. Class I peroxidases exist in the mitochondria, chloroplasts and bacteria. Class II peroxidases exist in fungi while Class III peroxidases are typical plant peroxidases (Welinder, 1992). Burel divided peroxidases into three types according to the isoelectric point: neutral (pI=7), alkaline or cationic (pI>7) and acid or anionic (pI<7) (Burel et al., 1994).

Peroxidases, as the important component of plant cells, have many physiological functions (Pandey et al., 2017; Passardi et al., 2005). They play an important role in the process of active oxygen metabolism (Inupakutika et al., 2016). Peroxidase can catalyze the NADH or NADPH in the cell wall to generate molecular oxygen and reactive oxygen species through a series of reactions (Montillet and Nicole, 2000). A large number of studies have showed that the activity of peroxidase is positively related to plant disease resistance (Almagro et al., 2009; Jwa and Hwang, 2017; Oliveira et al., 2017). The anionic peroxidase gene in tomato was reported to be induced by exogenous pathogenic fungi elicitor and H_2_O_2_ for the first time (Mohan and Kolattukudy, 1990). Expression of a tobacco anionic peroxidase gene was reported to be tissue-specific and developmentally regulated (Klotz et al., 1998). A highly anionic peroxidase was cloned in potato, with induced expression in suberizing potato tubers and tomato fruits. It was suggested to be involved in the deposition of the aromatic domain of suberin (Roberts et al., 1988). A few years later, the biochemical characterization of this suberization-associated anionic peroxidase of potato was reported (Bernards et al., 1999). However, few anionic peroxidase genes in potato were reported to be involved in late blight resistance except that RNA-silencing of anionic peroxidase gene M21334 was showed to decrease the potato plant resistance to *P. infestans* (Sorokan et al., 2014). More evidences of anionic peroxidase involved in potato disease resistance remain to be defined.

In our previous cDNA-AFLP study, a putative suberization-associated anionic peroxidase precursor (POPA) gene was found uniquely up-regulated by *P. infestans* (Li et al., 2009), which was further demonstrated by virus-induced gene silencing (VIGS) with TRV in *Nicotiana benthamiana* and potato to decrease the resistance to late blight (Du et al., 2013). Here we reported the cloning of this newly identified anionic peroxidase gene named *StPOPA* from potato and evidenced that the *StPOPA* gene enhanced potato resistance against *P. infestans* via promoting the accumulation of callose in cell wall and ROS in cytoplasm.

## Materials and methods

### Plant materials, pathogenic bacteria and plant treatments

*S. tuberosum* cultivar E-potato 3 (E3) was used in this study and transgenic E3 of 35S:GUS was used as empty vector control. Two physiological races of *P. infestans*, Ljx18 (1.3.4.7) and HB09-14-2 (1.2.3.4.5.6.7.8.9.10.11), collected from Hubei Province, China, were used to infect the transgenic and control plants.

To generate overexpression plants of the *StPOPA* gene, full-length cDNA was cloned using the primers (sequences of all the primers used in this study were listed in Supplementary Table S1) designed according to the PGSC sequence PGSC0003DMC400039701 from E3 and ligated to vector PBI121 via the restriction endonuclease site of *BamH*I and *Sac*I. To generate RNA-interference plants of the *StPOPA* gene, 170-bp non-conservative region of it and 308-bp conservative region of both the *StPOPA* gene and its homologous gene *StTAP2* were cloned with the primers and separately ligated to vector pHELLSGATE12 via recombination reaction. All the vectors mentioned above were transformed into *Agrobacterium tumefaciens* LBA4404.

For the biotic and abiotic stress treatments, leaflets with similar size on the third fully expanded leaves below the apical apex were excised carefully from the potato plants and sampled at 0, 6, 12, 24, 36, 48, 60 and 72 hours after infected by *P. infestans* and at 0, 3, 6, 9, 12, 24, 36, 48 and 60 hours after treated by SA (10 mM), ETH (100 μm), ABA (100 μm) and JA (50 μm). Sampled leaves were immediately frozen in liquid nitrogen and stored in −70 °C refrigerator till use.

### *Agrobacterium-mediated* genetic transformation of potato

*A. tumefaciens* LBA4404 containing target vectors (pBI121*-StPOPA* pHELLSGATE12-*StPOPA* and pHELLSGATE12-*StPOPA*-*TAP2*) were used to infect potato via microtuber discs transformation method (Tian et al., 2015). At first these agrobacterium strains were inoculated into 2 mL YEB liquid medium with 50 mg/L kanamycin (Kan) and 50 mg/L rifampicin (Rif), and the medium was cultured at 28 °C with 200 rpm rotation rate for 20 hours. Then 0.5 mL bacterial fluid was transferred into 50 mL fresh YEB liquid medium with 50 mg/L Kan, and the medium was cultured at 28 °C with 200 rpm rotation rate until the OD 600 reached 0.5.

*A. tumefaciens* bacterial fluid was centrifuged at 6,000 g at 4 °C for 6 minutes. Mycelium was then resuspended in equal volume of MS liquid medium (containing 3% sucrose). Seven to nine-week-old test-tube potatoes were cut into thin discs of 1–2 mm thick and these potato discs were soaked in *A. tumefaciens* bacterial fluid for 10 minutes. Positive lines were first screened on selective medium (MS + 3% sucrose + 1 mg/L indole-3-acetic acid (IAA) + 0.2 mg/L gibberellic acid (GA3) + 0.5 mg/L 6-benzylaminopurine (6-BA) + 2 mg/L zeatin (ZT) +75 mg/L Kan + 400 mg/L cefotaxime (Cef), PH 5.9) and then transferred to rooting medium (MS + 3% sucrose + 50 mg/L Kan + 200 mg/L Cef, PH 5.9), were then confirmed by the PCR with the primers of 35S promoter (the primers are shown in the Supplementary Table 1).

### *P. infestans* infection assays

*P. infestans* strain Ljx18 and HB09-14-2 (14-2) were cultured on rye and sucrose agar (RSA) medium at 18 °C for 13 days before collecting the inoculum for potato infection as previously described (He et al., 2015). Sporangia were collected from medium and washed with ddH_2_O to a concentration of 7×10^4^ per mL and then 0.01 mL sporangia suspension was used to inoculate potato leaves. Vernier caliper was used to measure the lesion length (L) and width (W). Lesion area (LA) was calculated by the formula LA=0.25×π×L×W. All the data were analyzed by ANOVA in the software SPSS.

### DNA, RNA extraction and real-time RT-PCR

Genomic DNA was extracted from potato leaves by the CTAB method and total RNA was extracted from potato leaves by the TRI pure Reagent (Aidlab). The first strand of cDNA was synthesized using M-MLV Reverse Transcriptase (Invitrogen) according to the manufacturer’s instructions. Quantitative PCR was conducted on a CFX connect™ Real-Time PCR Detection System (Bio-Rad) using SYBR Premix ExTaq (TaKaRa) according to the manufacturer’s instructions. The potato *EF-1α* gene was used as the internal control (primers were listed in Supplementary Table S1). The relative expression level was calculated as reported previously (Livak and Schmittgen, 2001).

### Transient expression in *Nicotiana benthamiana* and subcellular localization

Full-length CDS of the *StPOPA* gene was amplified with the adapters of attB1 and attB2, and then the target fragment was reorganized to target vector pK7FWG2, with GFP tag fusion expressed. Sequence confirmed vector was transformed into *A. tumefaciens* GV3101 and the single clone was cultured in 4 mL medium at 28°C with 200 rpm rotation rate. After 12 hours, bacterial fluid was harvested and resuspended in appropriate volume of MMA solution (10 mM MgCl_2_, 10 mM MES (Morpholine sulfonic acid), 200 μM AS (acetosyringone), pH 5.6) to adjust the OD_600_ of bacterial fluid to 0.01. Then, mix the target bacterial fluid with the p19 (gene silencing suppressor) (Voinnet et al., 2010) bacterial liquid with the volume ratio of 1:2. After 2 hours at room temperature, mixed bacterial liquid was injected into epidermis of *Nicotiana benthamiana.* After 36 hours of static growth, epidermis was torn off to be observed by fluorescence confocal microscope LSM510 (Carl Zeiss AG), GFP excitation light wavelength was 488 nm.

### Histochemical staining analysis

#### Trypan blue staining

Potato leaves inoculated with *P. infestances* for 5 days were soaked in Trypan blue solution (0.25 mg/mL) for 12 hours and then boiled to decolorize in the solution of lactophenol and absolute alcohol (volume ratio 1:2).

#### DAB staining

Potato leaves inoculated with *P. infestances* for 0, 6, 12 and 24 hours were soaked in the DAB (diaminobenzidine, Sigma) solution (1 mg/mL) for 8 hours and then boiled to decolorize as mentioned above.

#### Aniline blue staining

Potato leaves inoculated with *P. infestances* for 7 days were fixed in FAA solution (volume ratio of formaldehyde: acetic acid: absolute alcohol: water was 2:1:9:8), then leaves were soaked in 0.01% Aniline blue solution overnight and boiled to decolorize in solution of lactophenol and absolute alcohol (volume ratio 1:2). 0.02% fluorescent dye calcofluor was added onto decolorized leaves and samples were observed by positive microscope Zeiss Axioskop 40 (Carl Zeiss AG) in the ultraviolet excitation module (FT-395 nm, LP-42 nm). Callose was in yellow-green while sporangia and mycelium were in blue-purple.

### Preparation of transmission electron microscope samples

Potato leaves inoculated with *P. infestances* for 0, 12 and 24 hours were sampled for transmission electron microscope observation. The leaves around the inoculation site were cut into 0.1 cm × 0.2 cm pieces, and then those pieces were evacuated in 4% glutaraldehyde phosphate buffer (0.1 mol/L, PH 6.8). Vacuumized leaf samples were fixed in 4% glutaraldehyde for 3 hours, and then samples were rinsed by the phosphate buffer solution (pH6.8) for 5 times, 15 minutes each. Samples were then fixed in 1% osmic acid phosphate buffer (0.1 mol/L, PH 6.8) for 2 hours and rinsed by phosphoric acid buffer solution (pH6.8) for 4 times, 15 minutes each time. After that, the samples were dehydrated by 30%, 50%, 70%, 80%, 90% and 100% ethanol step by step, 30 minutes in each step. Then samples were dehydrated by 100% acetone for 3 times, 30 minutes each time before embedded with epoxy resin Epon812 and polymerized at 30 °C for 24 hours and 60 °C for 48 hours. Embedded sample blocks were positioned by glass cutter via semi thin section and positioned slice was made into ultra thin section by the diamond cutter. Slices were set onto the copper screen, and stained in 2% uranyl acetate for 10 minutes, following with staining in 2% lead citrate for 5 minutes. Slices were rinsed in distilled water and dried on a filter paper, and at last slices were observed and photographed via the transmission electron microscope JEM-1230 (JEOL).

## Results

### Isolation of the *StPOPA* gene in potato and cluster analysis of its homologous proteins

Cloning of the full-length cDNA of the *StPOPA* gene was done by electronic cloning (Du et al., 2013). The *StPOPA* gene has a length of 1092 bp encoding 363 amino acids. Cluster analysis of the *StPOPA* gene and its homologous genes in potato and other plants (data from NCBI database) was carried out, and it was indicated that three genes in tomato (gi|723674143, gi|225321568 and gi| 19359) had the highest similarity with the *StPOPA* gene (Fig 1A). The suberization-associated anionic peroxidase 2-like gene (gi|565360827), with the name *StTAP2* in the database, from potato clustered closely to the *StPOPA* gene was taken as a homologous gene of the *StPOPA* gene.

**Figure 1.**
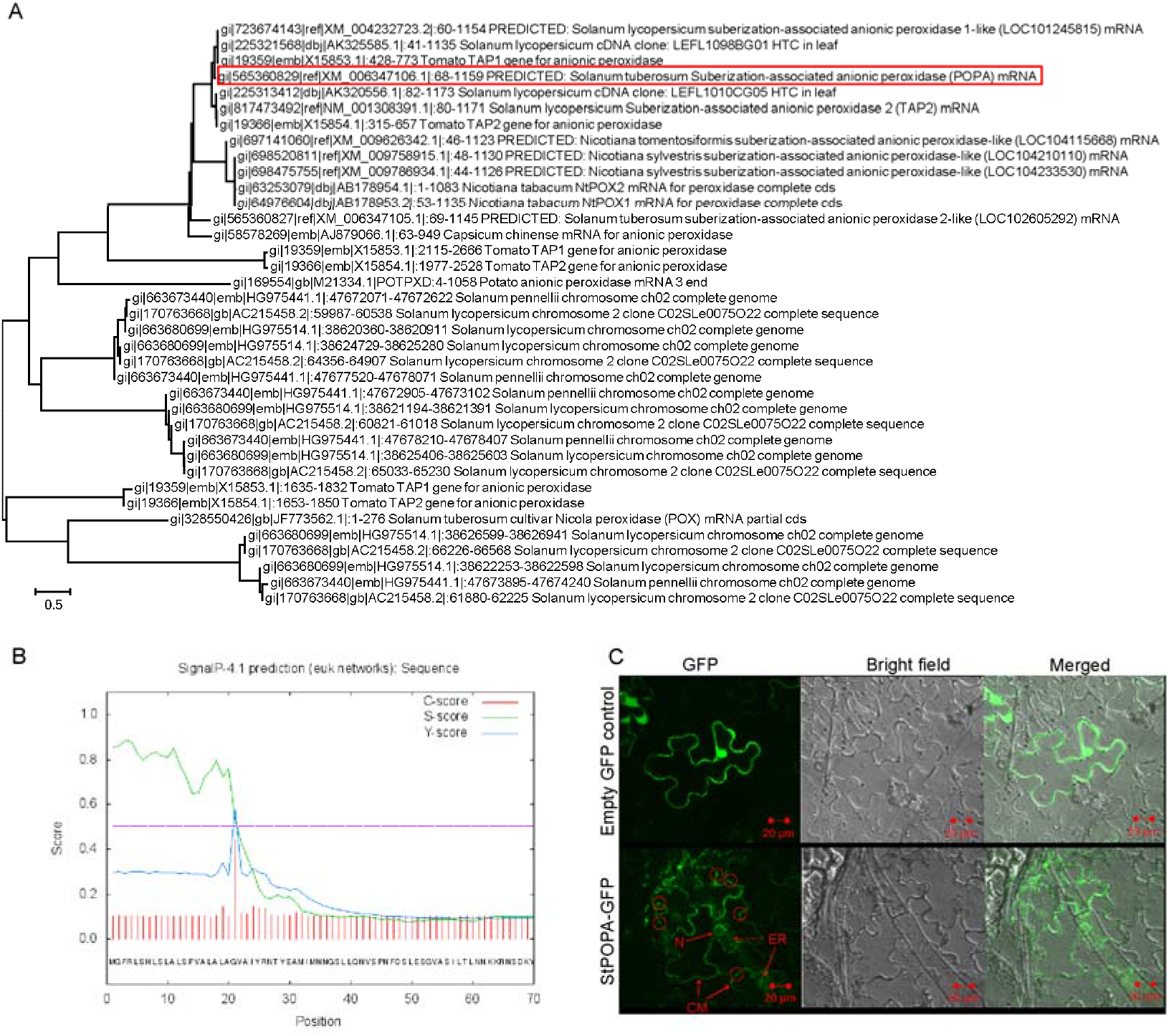
Cluster analysis, signal peptide prediction and subcellular localization of the StPOPA protein. A, Cluster analysis of the StPOPA homologous protein in potato and other plants. Red box represents the StPOPA clone in this study. B, Signal peptide prediction prediction of the StPOPA protein in SignalP 4.1 Server. C, Subcellular localization of the StPOPA protein in epidermal cells of *N. benthamiana.* Bright spots in red circles were possible secretory vesicles. CM, cell membrane. ER, endoplasmic reticulum. N, nucleus. Bar, 20 μM.

### Protein structure prediction and subcellular localization of the StPOPA

The signal peptide prediction result by SignalP 4.1 Server database (http://www.cbs.dtu.dk/services/SignalP/) showed that the StPOPA protein contained a significant signal peptide with the possibility of 0.790 (Fig 1B) and the subcellular localization prediction by the TargetP 1.1 Server database (http://www.cbs.dtu.dk/services/TargetP/) showed that the StPOPA protein might be located in the secretory pathway. To clarify the subcellular localization of the StPOPA protein in plant cells, we constructed a vector in which the StPOPA protein and GFP protein were fusion expressed. The subcellular localization of the StPOPA::GFP protein was observed by confocal after transforming *N. Benthamiana* for 36 hours. It was indicated that the StPOPA protein was mainly located in the cell membrane (CM), but we could vaguely observe GFP signal in the endoplasmic reticulum (ER), which was diffusely distributed around the nucleus. Moreover, we could even observe some small bright spots sporadically in the cell membrane, which were probably secretory vesicles (SV) (Fig 1C). These results suggested that the StPOPA protein was probably a secretory protein, and these results were very consistent with the characteristics of an enzyme.

### Expression profiling of *StPOPA* gene

In order to study the tissue expression profile and biotic and abiotic factors induced expression profile of the *StPOPA* gene, we detected the expression level of the *StPOPA* gene in different tissues of potato and the expression level of the *StPOPA* gene after *P. infestans* infection, mechanical damage and plant hormone treatment. It was shown that *StPOPA* gene could be detected in all the tissues with varied levels. The highest expression was found in the upper stems while the lowest in the tubers (Fig 2A).

**Figure 2.**
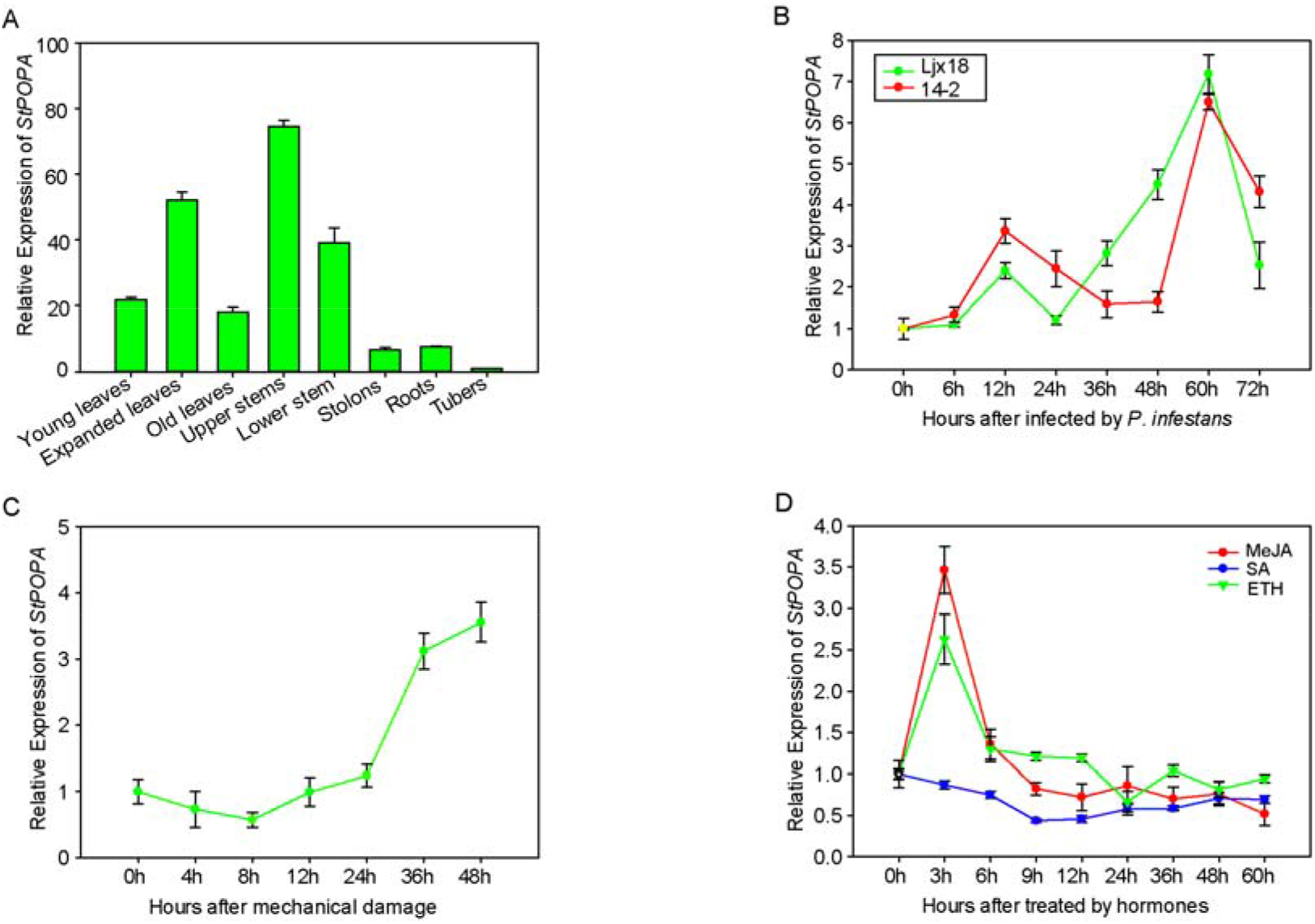
Tissue expression profile and induced expression profile of the *StPOPA* gene. A, Relative expression level of the *StPOPA* gene in representative tissues of potato. B, Relative expression level of the *StPOPA* gene in potato leaves after infected by *P. infestans.* C, Relative expression level of the *StPOPA* gene in potato leaves after mechanical damage. D, Relative expression level of the *StPOPA* gene in potato leaves after treated by hormones.

With regard to figure out whether the *StPOPA* gene was induced by the infection of *P. infestans*, we detected the expression level of the *StPOPA* gene in E3 after infection by two different *P. infestans* races, Ljx18 and 14-2 which performed incompatible and compatible interaction with E3, respectively. The results showed that the two races could both slightly induce the expression of *StPOPA* in the first 12 hours, followed by a rapid rise in transcriptional level at 24 hours inoculated with Ljx18 and at 48 hours with 14-2 while they both induced the highest expression of the *StPOPA* gene at 60 hours (Fig 2B). Our results demonstrated that *P. infestans* infection could induce an up-regulated expression of the *StPOPA* gene with an earlier response to incompatible race than compatible one, suggesting that the *StPOPA* gene may play a positive role against late blight in a relative late stage of the disease development.

The expression level of the *StPOPA* gene didn’t change significantly at the early stage after mechanical damage (Fig 2C), but the expression level of the *StPOPA* gene increased rapidly after 24 hours, implying that the *StPOPA* gene may have function in response to cell damage. Moreover, application of jasmonate (MeJA) and ethylene (ET) could activate the *StPOPA* gene to a high level in the first 3 hours but no significant effects were detected for salicylic acid (SA) (Fig 2D), speculating that the expression of the *StPOPA* gene could be regulated by the JA/ET signal pathway.

### Function dissection of *StPOPA* in resistance to *P*. *infestans*

In order to find out whether the *StPOPA* gene has a valid function in resistance to *P. infestans*, the function complementation test was conducted by overexpressing and silencing of the *StPOPA* gene, as well as simultaneously silencing *StPOPA* and its homologous gene *StTAP2* in potato variety E3. Selected transgenic lines of each transformation were illustrated in Fig. 3A-3C.

**Figure 3.**
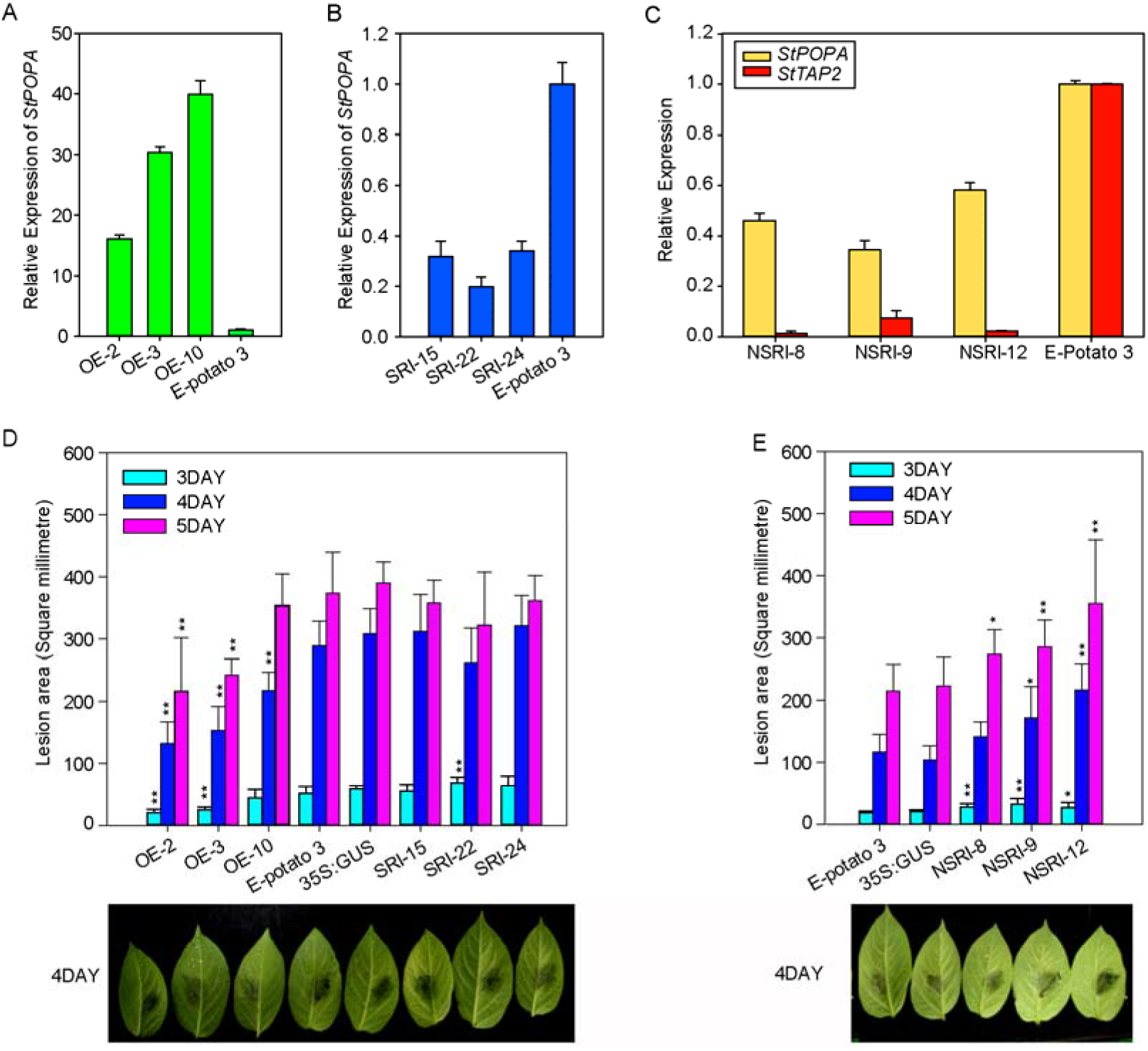
Resistance to late blight of the *StPOPA* gene overexpression and RNA-interference transgenic plants. A, Relative expression level of the *StPOPA* gene in leaves of overexpression plants. B, Relative expression level of the *StPOPA* gene in leaves of specific interference plants. C, Relative expression level of the *StPOPA* and *StTAP2* genes in leaves of nonspecific interference plants. D, Lesion area in leaves of the *StPOPA* gene overexpression and specific RNA-interference transgenic plants after *P. infestances* inoculation for 3, 4 and 5 days. Error bars indicate the standard deviation of 3 replicates. “*” indicates significant differences at p<0.05, “**” indicates extremely significant differences at p<0.01. E, Lesion area in leaves of nonspecific RNA-interference transgenic plants after *P. infestances* inoculation for 3, 4 and 5 days. Error bars indicate the standard deviation of 3 replicates. “*” indicates significant differences at p<0.05, “**” indicates extremely significant differences at p<0.01.

For evaluating the effects of the *StPOPA* gene on lesion expansion area, detached leaves of transgenic plants together with wild-type E3 and E3 transformed with the 35S:GUS vector were inoculated with *P. infestances.* It was indicated that leaf lesion area of overexpression lines was significantly smaller than wild-type and E3 transformed with 35S:GUS, whereas no significant difference was observed for the specific RNA-interference lines (Fig. 3D). However, silencing both the *StPOPA* and the *StTAP2* genes (nonspecific RNA-interference lines) exhibited a significantly larger lesion area than control (Fig. 3E), implying a redundant function of *StTAP2* to *StPOPA* in late blight resistance.

### Histochemical responses of transgenic leaves to *P. infestans* infection

The transgenic leaves together with control were stained using DAB to monitor the accumulation of H_2_O_2_ in cells. The H_2_O_2_ could be visualized after 6 hours of the inoculation. Overexpression of the *StPOPA* gene led to a higher accumulation of H_2_O_2_ while silencing the *StPOPA* gene resulted in a lower H_2_O_2_ abundance compared to wild-type and 35S:GUS transformed controls (Fig 4A). It was interesting that nonspecific interference of the *StPOPA* gene (i.e. silencing both *StPOPA* and its homologous *StTAP2*) had even a lower H_2_O_2_ production than the specific interference of the *StPOPA* gene. These results suggested that the *StPOPA* gene could transmit signal by reactive oxygen species to induce the resistance response to *P. infestances* infection.

**Figure 4.**
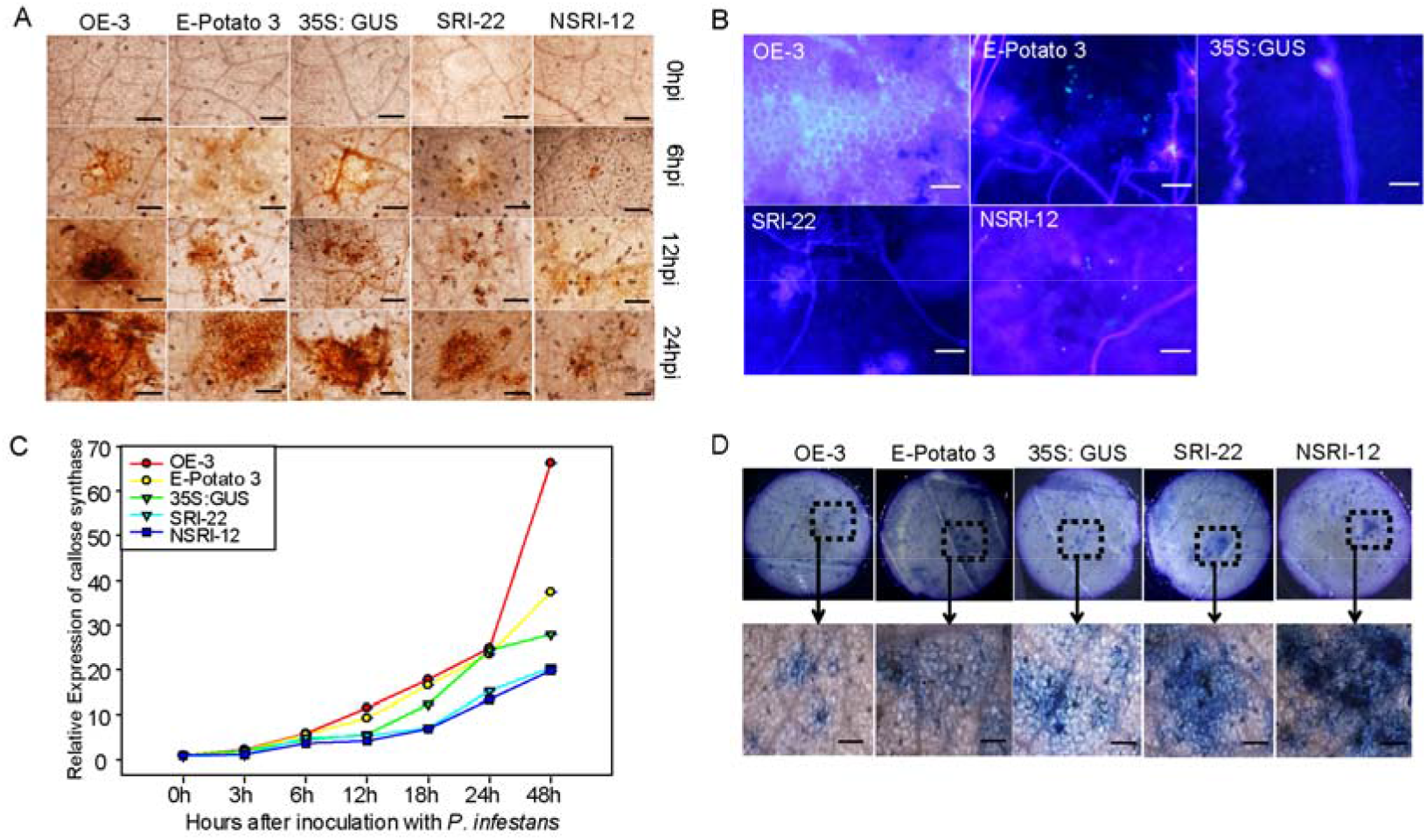
Accumulation of H_2_O_2_ and callose and cell death in leaves of the *StPOPA* gene overexpreesion (OE), specific silencing (SRI) and non-specific silencing (NSRI) plants. A, DAB staining of H_2_O_2_ for 0, 6, 12 and 24 hours after *P. infestans* inoculation. Bars, 200 μm. B, Aniline blue staining of callose 5 days after *P. infestans* inoculation. Bars, 50 μm. C, Relative Expression of β-1, 3 glucan synthase gene encoding the enzyme for biosynthesis of callose. D, Trypan blue staining of dead cells 36 hours after *P. infestans* inoculation. Bars, 200 μm. Wild type E-Potato 3 and 35S:GUS were taken as controls.

Similar results were observed in callose in the cells as stained by aniline blue. Abundant callose deposited around the point of inoculation in the leaves of overexpression lines and thickness of cell wall increased visibly (Fig 4B). On the contrary, no apparent accumulation of callose was observed in the leaves of specific interference and nonspecific interference lines. To confirm if the biosynthesis of callose was affected by the *StPOPA* gene, we tested the expression of β-1,3 glucan synthase, a key enzyme for callose synthesis. The full-length sequence of potato β-1,3 glucan synthase gene was obtained in NCBI database (*Solanum tuberosum* callose synthase 2-like, LOC102595703) and the expression level of β-1,3 glucan synthase gene in leaves of all the transgenic lines at multiple time points after *P. infestances* inoculation was detected by real-time quantitative PCR. From Fig 4C we could figure out that the relative expression level of β-1,3 glucan synthase gene increased during infection. In comparison to control, the relative expression level of the β-1,3 glucan synthase gene was remarkably elevated in overexpression line after 6 hours of *P. infestances* infection while the expression of the β-1,3 glucan synthase gene was significantly repressed in silenced lines. These findings implied that the *StPOPA* gene might play a key role in the process of callose formation by enhancing callose formation and deposit in the cell wall, withstanding the invasion and further spread of *P. infestance.*

The cell death around the inoculation point was detected by Trypan blue staining. Thirty-six hours after *P. infestances* inoculation, blue spots on the leaves of overexpression line were lighter than those of control but darker than control in the leaves of silenced lines with a more severe situation in the nonspecific interference line (Fig 4D). These results provided a conclusion that reinforcing the *StPOPA* gene could primarily reduce the cell damage by *P. infestans*.

### Effect of the *StPOPA* gene on cell ultra microstructure

With regard to elucidate the effects of the *StPOPA* gene on the pathogenesis of *P. infestances* in plant cells, the ultrastructure of the leaves sampled from each transgenic line after *P. infestances* inoculation was observed by electron microscopy (Fig 5). As a control, in the healthy potato leaves, chloroplasts in cells were regularly arranged on cytoplasmic membranes.

**Figure 5.**
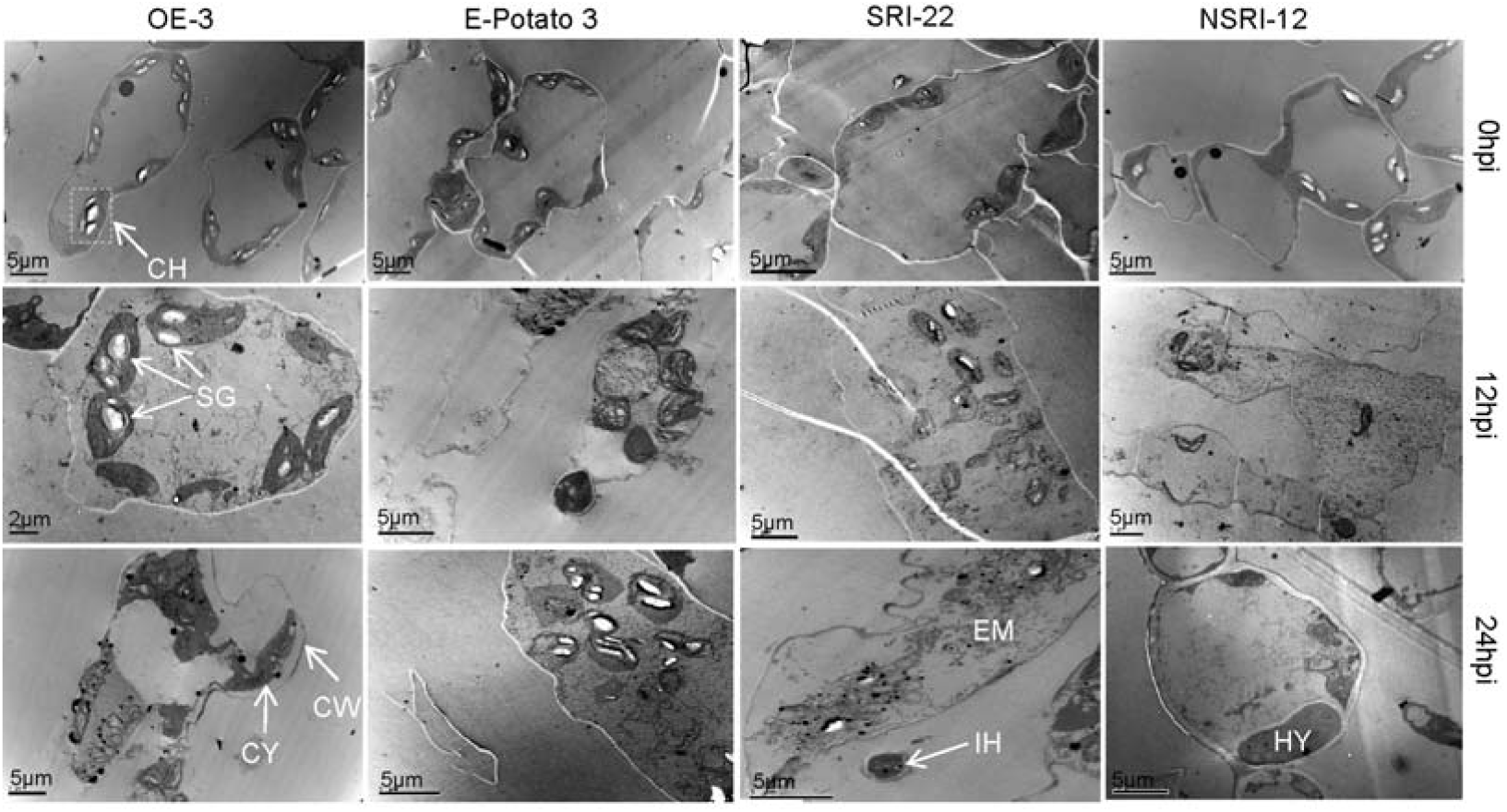
Transmission electron micrographs of leaves of the *StPOPA* gene overexpression (OE) and RNA-interference (SRI and NSRI) transgenic plants after *P. infestans* inoculation for 0, 12 and 24 hours. CH, chloroplast; CW, cell wall; CY, cytomembrane; EM, electron-dense material; HY, hyphae; IH, intercellular hyphae; SG, Starch granule.

There were no much variations in all leaf samples in the first 12 hours of *P. infestans* inoculation except that the mycelia of the pathogen extended along the cell wall of host cells or the gap between adjacent host cells in the leaves of overexpressed line. When the mycelia passed through cell wall of host cell, they formed haustellum between cell wall and cell membrane. After 24 hours, in the cells of leaves of the overexpressed line, plasmolysis was observed and much electron-dense material deposited in the space between the cell wall and the cell membrane. The outer membrane of chloroplasts disintegrated, the morphology of the chloroplasts gradually turned round and the laminated structure of thylakoid was slightly expanded. However, in the cells of leaves of specific interference line, the organelles of host cells such as the nucleus, chloroplast and endoplasmic reticulum were degraded while part of the complete organelle structure could still be observed. The more severe situation was in the cells of leaves of nonspecific interference line, the host cells were obviously necrotic, the organelles of adjacent cells were mixed together and completely degraded. These results were in accordance with the finding above that the *StPOPA* gene may have a function of maintaining cell integrity to reduce the damage caused by *P. infestans* infection.

## Discussion

Peroxidases are ubiquitous in plants (Milla et al., 2010) and they have a different degree of substrate specificity (Leon et al., 2002) and spatio-temporal expression pattern (Valério et al., 2004). The anionic peroxidase gene *StPOPA*, screened from our previous potato ESTs challenged with *P. infestans* (Du et al., 2013; Li et al., 2009), was proved to contribute to potato late blight resistance by complementary function dissection in the present research (Fig. 3). The *StPOPA* gene is constitutively expressed in potato plants, and its transcripts can be induced rapidly by jasmonic acid and ethylene and little latter by wounding and *P. infestans* (Fig. 2), suggesting that the *StPOPA* gene may have a function associated with cell damage in the signal pathway of JA/ETH.

It is noticeable that co-suppression of the *StPOPA* gene and its homologous gene *StTAP2* exhibited a higher resistance level to *P. infestans* than did by suppressing only the *StPOPA* gene (Fig. 3D). Previous study showed that biotic and abiotic stress could both induce the expression of different peroxidase isoenzymes (Navrot et al., 2006). Besides, there were reports also identified the similar function of different perioxidase isoforms (Reumann and Bartel, 2016). Therefore, we speculated that difference in late blight resistance of specific interference of the *StPOPA* gene and nonspecific interference of both the *StPOPA* and *StTAP2* genes may come from the functional redundancy of these homologous genes.

It is well understand that oxidative burst characterized by accumulation H_2_O_2_ in plant cells is one of the defense responses of plants against pathogen infection. Plants usually generate reactive oxygen species (ROS) as signaling molecules that activate various processes including pathogen defense, programmed cell death, and stomatal behavior (Apel and Hirt, 2004). In the present research, we clarified that H_2_O_2_ was predominantly formed by overexpressing the *StPOPA* gene (Fig 4A). A possible quantitative correlation between the *StPOPA* gene expression and H_2_O_2_ abundance was observed. A possible explanation could be that, as a peroxidase, StPOPA catalyzes the formation of H_2_O_2_ to control the process of plant defense against the invasion of *P. infestance.* It was indicated that in the process of pathogen defense response, the plant simultaneously produced more ROS while decreasing its ROS scavenging capacities, this led to the accumulation of ROS and activation of programmed cell death (PCD) (Delledonne et al., 2001; Foyer and Noctor, 2000). Our results demonstrated that severe cell death was accompanied with the suppression of the *StPOPA* gene, indicating a potential mode of the *StPOPA* gene to defeat late blight by enhancing H_2_O_2_ production for purposeful programmed cell death.

It was documented that, in the presence of H_2_O_2_ and phenolic substrates, peroxidases could act in the peroxidatic cycle and were engaged in the synthesis of lignin and other phenolic polymers (Apel and Hirt, 2004). When the pathogen infects, the cell wall rehabilitation and reinforcement are the first physiological barrier to resist the pathogen invasion. The cell wall would accumulate a lot of callose and lignin for cell wall lignification and the formation of the mastoid. In our study, overexpression of the *StPOPA* gene could promote the formation of callose and possibly other cell wall sediments and the deposition of electron-dense materials (Fig 5). This finding was further confirmed by a higher expression level of the −1,3 glucan synthase gene in the *StPOPA* gene overexpressed plants than control and the *StPOPA* gene silenced plants (Fig. 4C). Synthesis of callose possibly by enhancing β-1,3 glucan synthase was thus considered an effective way for the *StPOPA* gene to defense against the infection of *P. infestans.* This process led to a significant inhibition of the pathogen growth and expansion in the host cells of the *StPOPA* gene overexpressed plants (Fig. 4D).

And a little bit more, our results showed that overexpression of the *StPOPA* gene was correlated with a decrease in the resistance and a marked increase in ROS and callose synthesis, so we could conclude that overexpression of the *StPOPA* gene increased potato resistance to *P. infestans.* It was reported that the apolastic oxidative burst generated by peroxidases was an important component of PTI and it could contribute to DAMP-elicited defenses such as callose synthesis (Daudi et al., 2012). However, overexpression of anionic peroxidases was known to affect plant development in some cases, especially root growth (Lagrimini et al., 1997). So there was a possibility that the generation of ROS in the *StPOPA* overexpression lines might just a consequence of the deregulation of the apoplastic peroxidases synthesis. And it’s highly likely that the primary role of anionic peroxidases in the resistance to pathogens, was to use rapid hydrogen peroxide to cross-link the different phenolic monomers occurring in lignin or in the phenolic domain of suberin (Quiroga et al., 2000). In order to clarify the exact mechanism of the *StPOPA* gene, more work remained to be done in our future research.

In conclusion, our present research revealed that the *StPOPA* gene contributed to potato resistance against *P. infestans.* On the one hand, the *StPOPA* gene could promote H_2_O_2_ burst and cell necrosis, restraining the infection of pathogenic bacteria and expansion of mycelium. And on the other hand, enhanced peroxidase activity may catalyze the synthesis of callose, thus preventing the invasion of pathogenic bacteria and its expansion in the host cells.

## Acknowledgments

This research was supported by grants from the National Natural Science Foundation of China (31261140362).

## Supplementary data

Supplementary Table S1. Primers used in this study.

